# Epidermal and ECM Damage Following Pinch Injury Restricts Dendrite Regeneration in *Drosophila*

**DOI:** 10.64898/2026.07.15.738747

**Authors:** Mia A Brantley, Avantika Pandiyan, Annie C Danh, Sydney E Prange, Dario S Rimicci, Katherine L Thompson-Peer

**Author notes:** Corresponding author: Correspondence addressed to: Katherine Thompson-Peer. MB designed research, performed research, analyzed data, and wrote the paper. AP analyzed data. AD analyzed data. SP performed research. DR performed research. KTP designed research, analyzed data, and wrote the paper.

## Abstract

Neuronal dendrites can be injured by a number of insults, but the cellular mechanism by which dendrites respond to tissue injury and undergo repair is poorly understood. Much of the field’s progress has evaluated dendrite regeneration following laser injury. While precise, laser injury does not accurately model the real-world damage to surrounding tissue that would accompany neuronal injury. Here, we modify a pinch injury technique to injure both the dendrites and their surrounding tissues in *Drosophila melanogaster* larvae, more similar to what is observed in real-world injury. We refined this technique such that only half of a sensory neuron’s dendrites are injured, leaving the other half uninjured. Our data indicate that both dynamic and stable dendritic arbors regrow dendrites following pinch injury. Neurons primarily engage in compensatory regeneration whereby new branches are added on the uninjured half of the arbor. Comparing the regenerative response following pinch versus laser injury revealed that dendrites preferentially regrew into areas where the surrounding tissue was left intact, and not into areas where the surrounding tissue was damaged by pinch. These results prompted us to evaluate the damage sustained to surrounding tissue. In examining non-neuronal tissues after pinch injury, we found damage to epidermal cells and the ECM, but not glia. We also observed a robust immune response on the pinched half of the arbor. We conclude that the sustained damage to surrounding tissue and the initiation of an immune response create a non-permissive environment for dendrite regeneration following pinch injury.

**Significance Statement:** Neuronal dendrites are injured in clinical conditions, such as stroke, traumatic brain injury, and neonatal hypoxia. Dendrites also degenerate in the early stages of a number of neurodegenerative diseases. The role of surrounding tissues in dendrite regeneration is poorly characterized, especially considering that neuronal injury is typically accompanied by broad tissue damage. Our data evaluates dendrite regeneration following an injury that better mirrors real-world conditions and demonstrates that broad tissue damage diminishes a neuron’s capacity to regenerate its dendrites. Our findings show that neurons preferentially regrow into intact, undamaged tissue environments, addressing a large gap in the field’s knowledge: how damage to the surrounding tissue limits neuron regeneration after injury.

**Visual Abstract:** 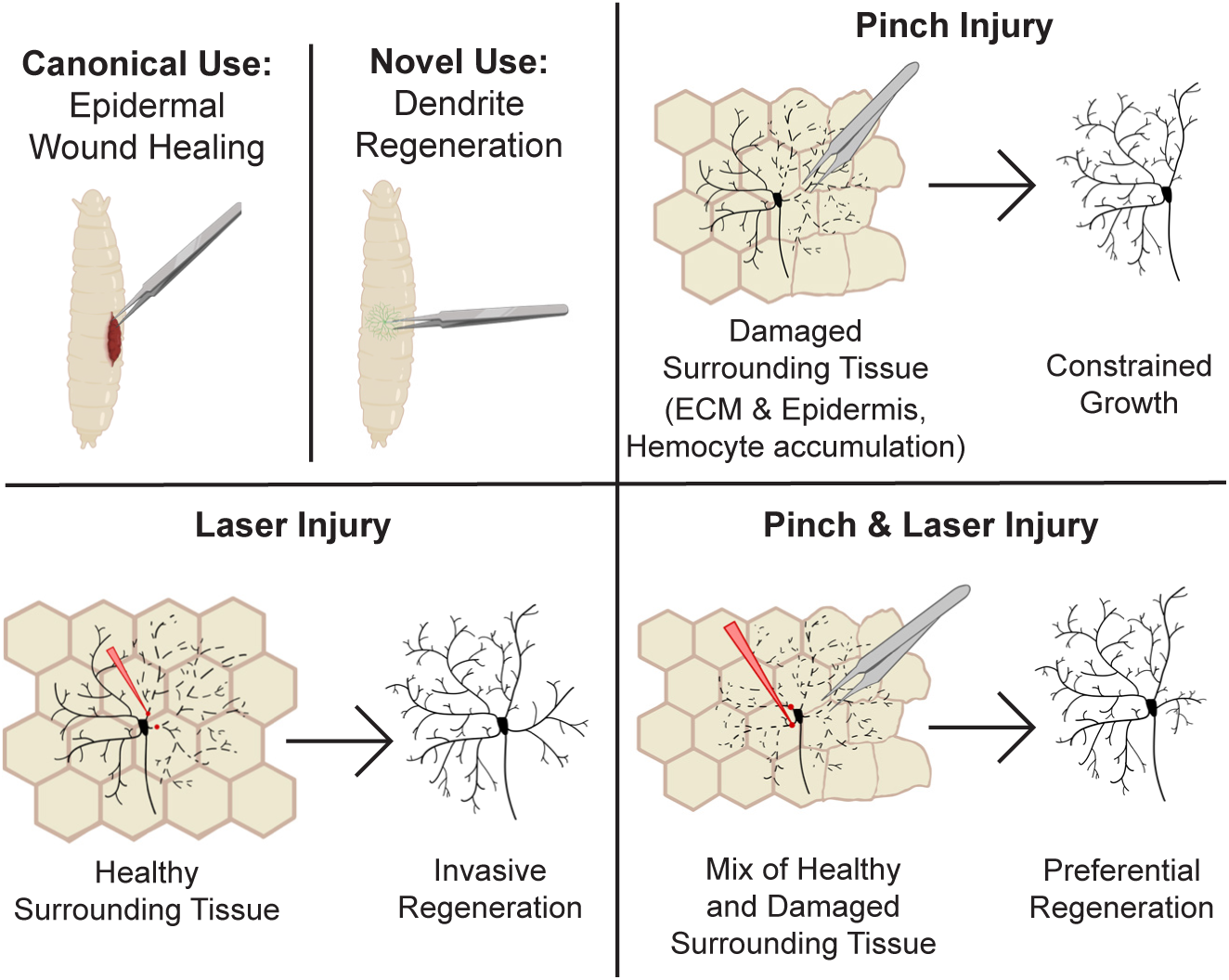

## Introduction

Neurons are the main signal transducers of the nervous system. They are composed of two distinct structures: dendrites that receive signals, and axons that send those signals. Unfortunately, dendrites can be injured by a number of insults like stroke, hypoxia, repeated traumatic brain injuries (TBI), and neurodegenerative disease. In response to this damage, dendrites are capable of regeneration. Following the removal of all dendrite branches by laser injury, dendrites of the *Drosophila melanogaster* peripheral nervous system (PNS) can regenerate the same number of branches as age-matched uninjured control neurons (Song et al., 2012; Stone et al., 2014; Thompson-Peer et al., 2016). Similarly, PVD neurons of the *Caenorhabditis elegans* PNS can regenerate their dendrites via plasma membrane fusion (Oren-Suissa et al., 2017; Brar et al., 2022). Motor neurons in the zebrafish spinal cord regenerate their dendrites 5-7 days post laser microdissection, demonstrating that dendrites of the central nervous system (CNS) are also capable of regeneration (Stone et al., 2022). While dendrite regeneration in the mammalian CNS is limited, it has been observed immediately following laser microsurgery in the mouse brain and spinal cord (Zhao et al., 2017). Further, dendrite regeneration in the adult mouse cerebral cortex has been observed following brain prick injury (Paveliev et al., 2016).

To investigate the mechanisms of dendrite regeneration, we use the dendritic arborization (da) sensory neurons of the *Drosophila* PNS (Grueber et al., 2002). These da neurons are grouped by class (I-IV) and innervate the epidermis of the larval body wall. The development of the da neural system is heavily dependent upon a permissive extracellular environment and signals from surrounding cell types and tissues. For example, an epidermally-derived microRNA, *bantam*, regulates and restricts the scaling growth of class IV da neurons (Parrish et al., 2009; Jiang et al., 2014). Further, proper space-filling of class IV da neurons requires a permissive signal created by the presence of Heparan Sulfate Proteoglycans (HSPGs) on the epidermal cell surface (Poe et al., 2017). The extracellular matrix (ECM) also plays a critical role in dendrite development in other systems (Dansie and Ethell, 2011; Levy et al., 2014; Long and Huttner, 2019; Heiman and Bülow, 2024).

In addition to normal development, recent studies hint that dendrite regeneration of da neurons after injury is likely regulated by extracellular signals. Epidermally-derived *bantam* signaling restricts dendrite regeneration in class IV ddaC neurons (Song et al., 2012). Regenerated dendrites exhibit a decreased adherence to the ECM, which is mediated by integrins, resulting in a failure of self-avoidance mechanisms (Thompson-Peer et al., 2016). Devault and colleagues (2018) revealed that dendrite regeneration in adult flies is constrained by ECM remodeling as a result of epidermally-expressed matrix metalloproteinase 2 activity (DeVault et al., 2018). Alterations in attachment to the ECM may result in many of the morphological differences that newly regenerated dendrites display, such as decreased area coverage (Thompson-Peer et al., 2016).

Dendrite regeneration is a novel and growing field. In comparison to axon regeneration, dendrite regeneration has not been extensively studied (Peterson and Benowitz, 2018). One reason for this is because traditional surgical injury approaches are not suitable for dendrite injury (Varier et al., 2022; Hertzler and Rolls, 2024; Vaughn and Lee, 2024). Recent advancements in microscopy have overcome these challenges to facilitate studying dendrite regeneration (Song et al., 2012; Stone et al., 2014; Thompson-Peer et al., 2016; DeVault et al., 2018; Nye et al., 2020; Hertzler et al., 2023; Duarte et al., 2024; Hertzler and Rolls, 2024; Prange et al., 2024; Hwu et al., 2025). However, these studies utilized a laser injury method, which is slow, has a low throughput, and does not accurately model real-world neuronal damage. The lack of real-world modeling is especially important given that the role that extracellular cues play in dendrite regeneration needs deeper discovery.

A means of more accurately modeling real-world injury has been demonstrated by the development of an epithelial pinch wound assay to study wound healing and reepithelization in *Drosophila* larvae (Galko and Krasnow, 2004; Stevens and Page-McCaw, 2012; Burra et al., 2013; Tsai et al., 2017). This pinch wounding assay has also been utilized for an RNAi screen to identify genes involved in proper wound healing (Lesch et al., 2010). It has additionally been adapted to evaluate ECM wound healing (Ramos-Lewis et al., 2018).

Here, we have established a pinch injury technique to injure the da neurons and surrounding tissues. We found that this pinch injury damaged class IV and class I da neurons, the epidermis, and the ECM, while leaving the glia largely uninjured. This injury method also stimulated invasion of hemocytes into the injured area. Response to this damage was stereotyped: dendrites preferentially regrew into empty territory where the surrounding tissue was left intact, while surrounding tissue damage restricted dendrite regeneration. This pinch injury method was faster and more accurately modeled damage to the neuron, its dendrites, and surrounding tissue. Overall, we find that damage to the ECM and other surrounding tissues impedes dendrite regeneration.

## Methods and Materials

### Experimental Model and Subject Details

#### Drosophila strains

Drosophila stocks were maintained at room temperature. The following fly strains were used in this study: Canton-S, ppk-CD4-tdGFP (second chromosome, BDSC #35842), ppk-CD4-tdGFP (third chromosome) (Han et al., 2011), Gal4^ppk^ (second chromosome) (Grueber et al., 2003), UAS-CaMPARI2.L389T (second chromosome, BDSC #78319) (Moeyaert et al., 2018), Gal4^2-21^, UAS-CD4-tdTomato (third chromosome), ppk-CD4-tdTomato (second chromosome, BDSC #35844), Gal4^A58^ (third chromosome) (Galko and Krasnow, 2004), viking-GFP Protein Trap (second chromosome, BDSC #98343), UAS-viking-GFP (third chromosome, courtesy of Noselli Lab) (Van De Bor et al., 2015), UAS-Armadillo-GFP (third chromosome, BDSC #58725), Gal4^repo^, UAS-mRFP (third chromosome) (Sepp et al., 2001), Gal4^pcn^ (third chromosome, BDSC #600223), UAS-CD4-tdTomato (second chromosome, BDSC #35841).

A complete list of fly stocks can be found below in the Key Resources Table.

### Generation of Fly Lines and Experimental Crosses

Crosses were performed in an incubator at 22.5°C and 70% humidity and eggs were collected on a plate made of grape juice and agarose (grape plate) with yeast paste to synchronize animal age. Cross progeny larvae for experiments were kept on grape plates in an incubator at 22.5°C and 70% humidity until used for injuries. After injury, larvae were individually housed in grape plates with yeast paste and maintained at 20°C and 70% humidity until the last imaging time point (72 hours after injury). Both male and female larvae were used for all experiments. Assays to assess pinch or 2p injury and subsequent regeneration in class IV ddaC neurons were performed by crossing fly lines expressing ppk-CD4-tdGFP with Canton-S flies. Assays to assess pinch injury and subsequent regeneration in class I ddaE neurons were performed by crossing fly lines expressing Gal4^2-21^ and UAS-tdTomato with Canton-S flies. Assays to assess damage to whole-body ECM were performed by crossing flies expressing ppk-CD4-tdTomato to flies expressing the viking-GFP protein trap. Assays to assess damage to A58-driven ECM were performed by crossing flies expressing ppk-CD4-tdTomato and Gal4^A58^ to flies expressing UAS-viking-GFP. Assays to assess damage to epidermal cells were performed by crossing flies expressing ppk-CD4-tdTomato and Gal4^A58^ to flies expressing UAS-Armadillo-GFP. Assays to assess damage to glia were performed by crossing flies expressing Gal4^repo^ and UAS-mRFP with flies expressing ppk-CD4-tdGFP. Assays to assess the immune response following injury were performed by crossing flies expressing Gal4^pcn^ and UAS-CD4-tdTomato with flies expressing ppk-CD4-tdGFP. Assays to assess CaMPARI conversion were performed by crossing flies expressing UAS-CaMPARI2.L389T with flies expressing ppk-Gal4^vk34^ and ppk-CD4-tdGFP.

### Dendrite Injury Assays

All injury assays were performed in live, whole-mount larvae.

### Pinch Injury Injury Assay

Please refer to the schematic in Fig. 1A for a visual explanation of the experimental flow-through of the pinch injury assay. Please refer to multimedia file 1 for a video recording of a pinch injury. To perform the pinch injury assay, groups of 5-6 age-matched 96 hours after egg lay (AEL) third instar larvae were collected onto a pad of 4% agarose, and the pad was transferred to a small, empty petri dish. The small petri dish was placed into a larger petri dish for anaesthetization. Larvae were anaesthetized with isoflurane for 45-50 seconds by placing a folded-up Kimwipe moistened with a small amount of isoflurane in the larger petri dish, exposing the larvae to the isoflurane vapor. After anaesthetization, the larvae, sitting on the agarose pad, were transferred out of the petri dish and visualized under a light dissecting microscope. Larval segments (thoracic and abdominal) were counted until reaching abdominal segments 4 and 5. Larvae were rolled ∼30° to expose the right half of the dorsal side of the animal for pinch injury. A pair of blunted #5 forceps were used to pinch the animal. One tine of the forceps was aligned with the anterior side of abdominal hemisegment 4 (A4); one tine of the forceps was aligned with the posterior side of abdominal hemisegment 5 (A5). A slight scooping motion was used to properly pick up and pinch the two hemisegments together: the anterior side of A4 meets the posterior side of A5, with the cuticle region between the forceps popping out, resembling a “muffin top.” (Please see Figure 1A for a cross-sectional reference of the larval “muffin top.”) The pinch was held with minimal pressure for 10 seconds and then released. For CaMPARI experiments, larvae were exposed to 20 seconds of UV light immediately after pinching. Individual larvae were then housed in grape plates with yeast paste and maintained at 20°C and 70% humidity.

**Figure 1:**
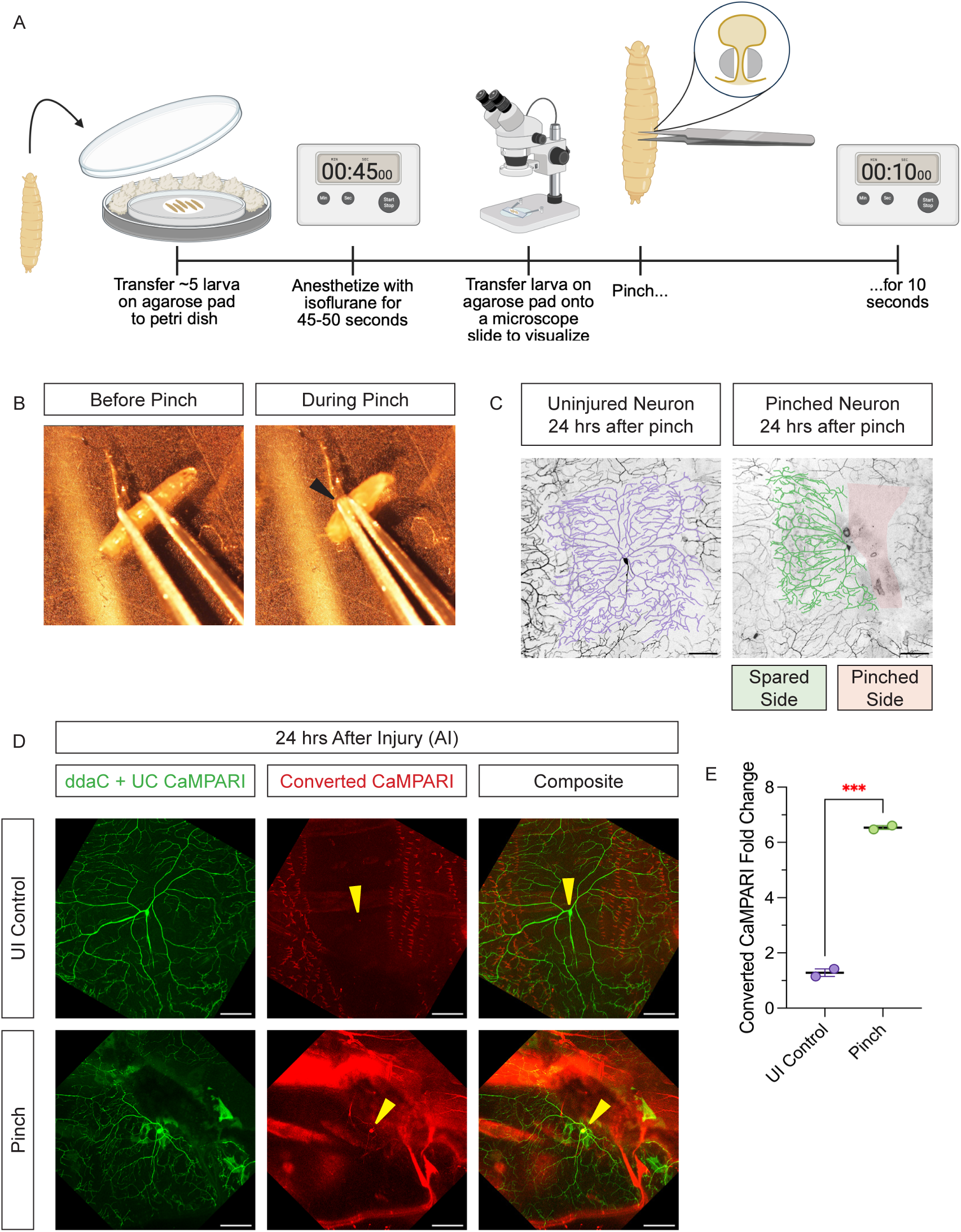
Pinch Injury Damages Peripheral Sensory Neurons in *Drosophila* Larvae **(A)** Schematic of pinch injury technique. **(B)** Larvae before and during pinch injury. **(C)** An uninjured (UI) class IV ddaC neuron and a pinched injured neuron 24 hrs after pinch injury. Pinch injury damages half of the dendritic arbor, leaving the other half spared. Scale bar 100 µm. **(D)** Uninjured (UI) within-animal control neurons and pinched neurons 24 hrs after injury (AI), expressing both CD4-tdGFP and CaMPARI, which photoconverts from green (unconverted (UC) CaMPARI) to red (converted CaMPARI). Yellow arrowhead indicates cell body. Scale bar 100 µm. **(E)** Red converted CaMPARI fluorescence (normalized to background fluorescence) in uninjured (UI) within-animal control neurons (purple) and pinched neurons (green) at 24 hrs AI.

### Two-Photon Injury Assay

Larvae were immobilized for mounting by being sandwiched between an agarose pad and coverslip held together with vacuum grease (Thompson-Peer et al., 2016; Duarte et al., 2024; Prange et al., 2024; Hwu et al., 2025). Glycerol was used as the mounting media. Two-photon (2p) injury assays were performed with the Spectra-Physics MaiTai two-photon tunable laser mounted on the Zeiss LSM980 confocal microscope. Neurons were imaged using 488 nm green or 561 nm red lasers and injured using the bleaching function with the 2p laser at 820 or 860 nm. Exposure to the laser lasted for ∼1.5 s. For the balding injury assay, the laser was focused on primary dendrite branches near the cell body to remove all branches with the fewest cuts possible without damaging the cell body. For the half-2p injury assay, the laser was focused on primary or secondary dendrite branches to remove half of the branches in the dendritic arbor with the fewest cuts possible. After laser injury, individual larvae were housed in grape plates with yeast paste and maintained at 20°C and 70% humidity. Successful balding injury was assessed by observing dendrite blebbing immediately after injury, comparing the 24 hours AI images to the before injury images to assess if the arbors look different, and evaluating territory coverage of new arbors at 24 hours AI.

### Half-Pinch + Half-2p Injury Assay

Larvae were pinched according to the details listed above in “Pinch Injury Assay.” Two hours later, larvae were mounted as detailed above and the pinch injured neuron was subjected to 2p laser injury. By two hours after pinch injury, the blebbing, degradation, and removal of damaged branches was observed before reliably performing a secondary 2p laser injury. All remaining branches were removed. After laser injury, individual larvae were housed in grape plates with yeast paste and maintained at 20°C and 70% humidity.

### Imaging

Larvae were immobilized for mounting by being sandwiched between an agarose pad and coverslip held together with vacuum grease (Thompson-Peer et al., 2016; Duarte et al., 2024; Prange et al., 2024; Hwu et al., 2025). Glycerol was used as the mounting media. Images of injured and within-animal uninjured control neurons were taken on a Zeiss LSM700 confocal microscope. Neurons were imaged at 24 hrs AI to confirm injury. Neurons were imaged at 72 hrs AI to observe the end time point of regeneration. Only neurons with obvious survival and absence of half the dendritic arbor were included in subsequent analyses and presented data.

### Quantification and Statistical Analysis

Dendrite arbors were traced using the Simple Neurite Tracer (SNT) plugin in ImageJ to determine the number of dendrite branch tips and the total length of all the dendrite branches (Schindelin et al., 2012). The polygon selection tool in ImageJ was used to select and measure the total area of the dendritic arbor. For all graphs comparing means, values are plotted as mean ± standard error of the mean (SEM). For all graphs comparing means, individual data points represent individual neurons, except for Figure 1-1A, in which individual data points represent the average time spent performing the pinch or half-2p injury assay. For paired data (Fig 2B, Fig 2C, Fig 2F, Fig 2G, Fig 3 - 1B, Fig 3-2C), values are plotted for individual neurons with faded lines connecting repeated measurements of the same neuron at 24 and 72 hrs AI. Solid lines represent the slope (m) between the mean 24 and 72 hrs AI values, which was calculated using the following equation:

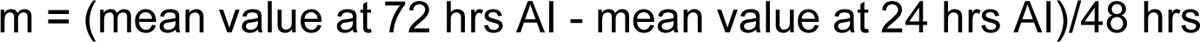

**Figure 2:**
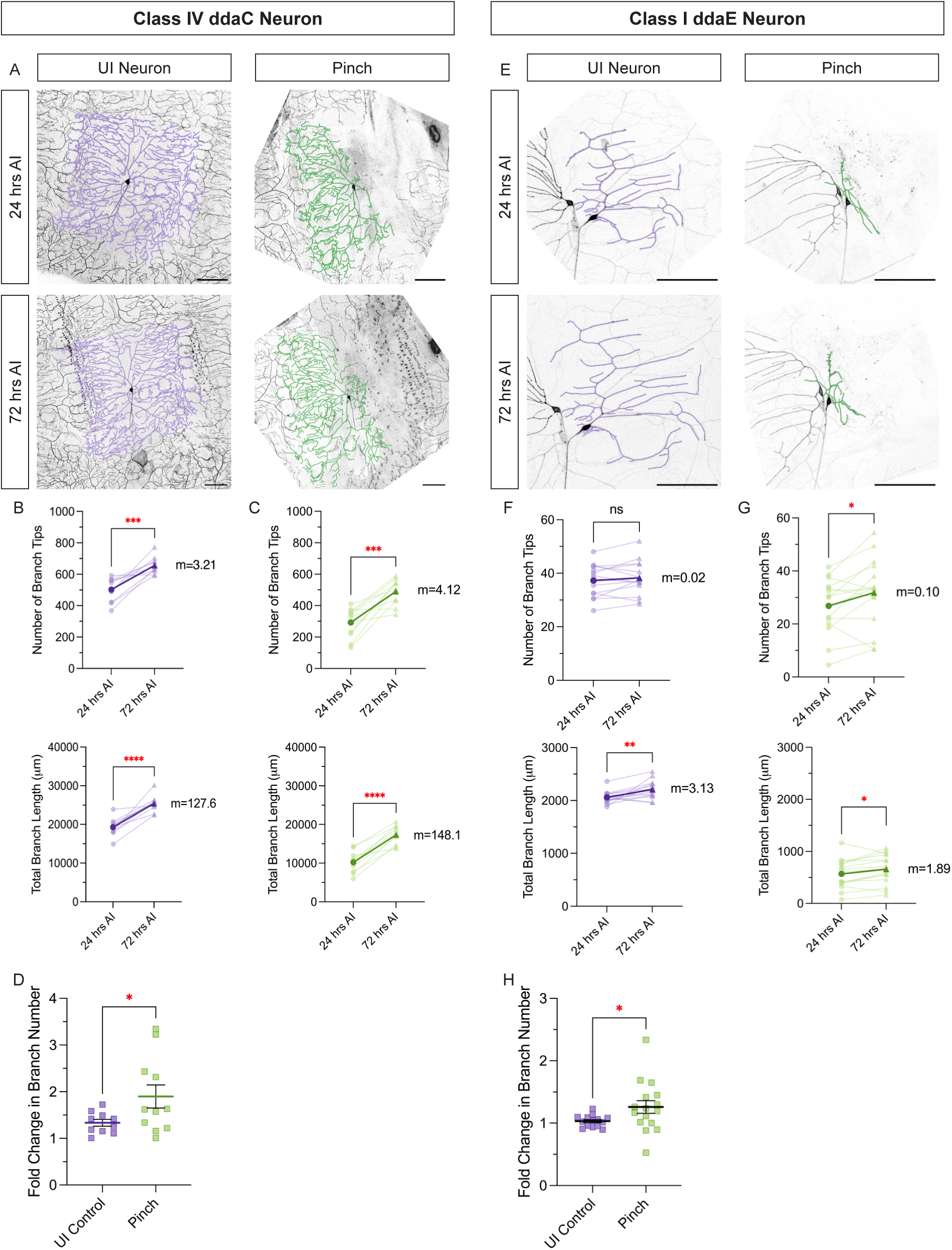
ddaC and ddaE Neurons Grow Following Pinch Injury. **(A)** Class IV ddaC uninjured (UI) within-animal control (purple) and pinched (green) neurons at 24 and 72 hrs AI. Scale bar 100 µm. **(B-C)** Number of branch tips (top) and total branch length (bottom) of UI neurons (B) and pinch injured class IV ddaC neurons (C) at 24 and 72 hrs AI. **(D)** Fold change in branch number of UI and pinched class IV ddaC neurons. **(E)** Same as A, but class I ddaE neurons. Scale bar 100 µm. **(F-G)** Same as B-C, but class I ddaE neurons. **(H)** Same as D, but class I ddaE neurons.

**Figure 3:**
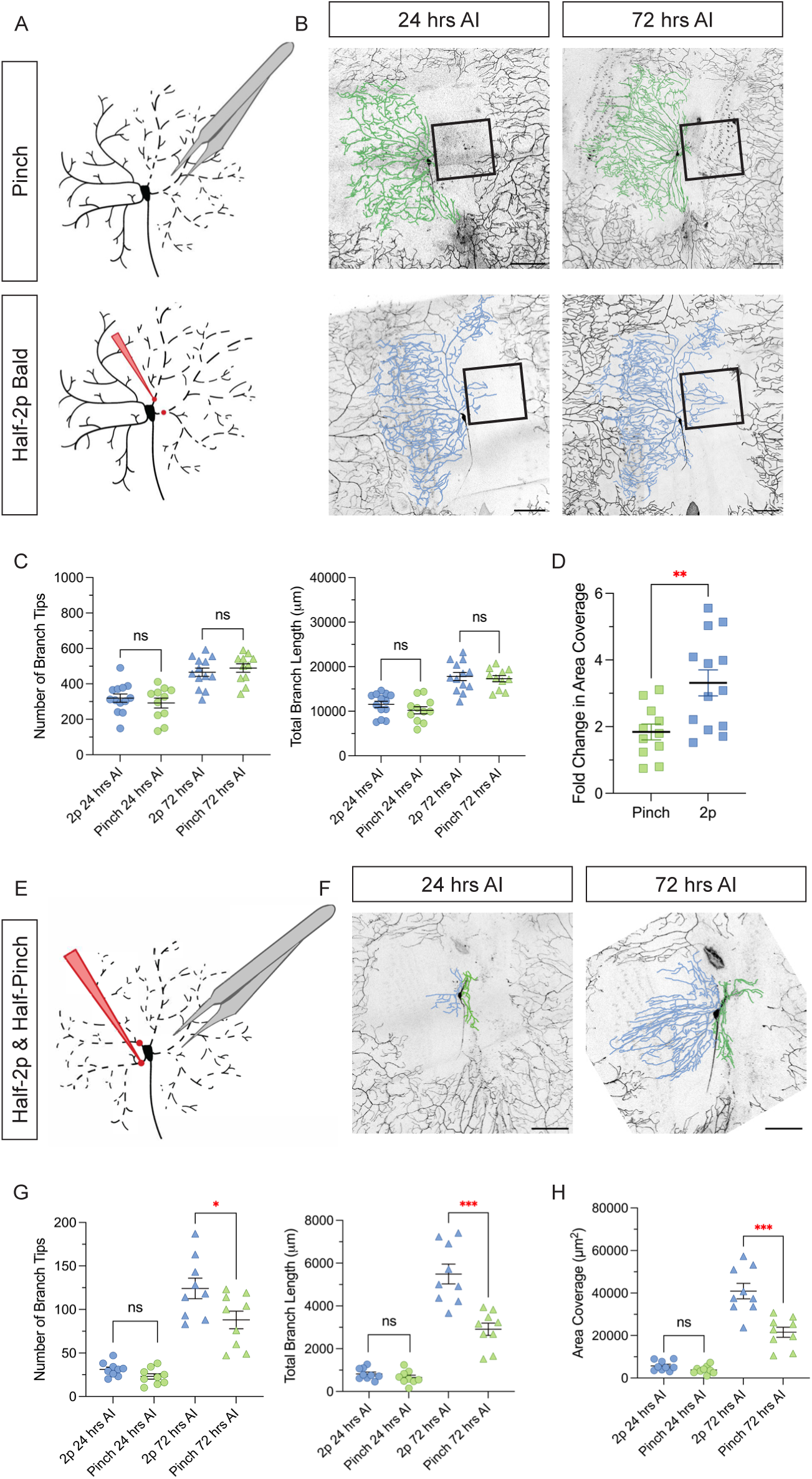
ddaC Neurons Preferentially Regenerate Into Uninjured Territory. **(A)** Schematics of pinch (top) and half-2p bald (bottom) injuries. **(B)** Pinch (top) and half-2p bald (bottom) injured class IV ddaC neurons at 24 and 72 hrs AI. Black boxes identify empty territory. Scale bar 100 µm. **(C)** Number of branch tips (left) and total branch length (right) of class IV ddaC neurons injured by 2p (blue) or pinch (green). **(D)** Fold change in area coverage of the injured half of individual neurons following 2p or pinch injury. **(E)** Schematic of a “half-pinch + half-2p” injury on the same class IV ddaC neuron. **(F)** Half-2p (left side) + half-pinch (right side) injured class IV ddaC neuron at 24 and 72 hrs AI. Scale bar 100 µm. **(G)** Number of branch tips (left) and total branch length (right) of 2p injured side (blue) and pinch injured side (green) of individual class IV ddaC neurons injured by both injury paradigms. **(H)** Area coverage of 2p injured side (blue) and pinch injured side (green) of individual class IV ddaC neurons injured by both injury paradigms.

Fold change in branch number was calculated using the following equation:

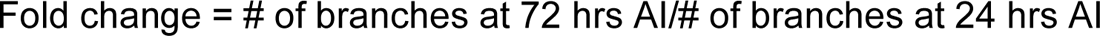

To calculate area coverage, we used the axon as a midpoint guideline for which branches to include as part of the injured half of the arbor. Fold change in area coverage was calculated using the following equation:

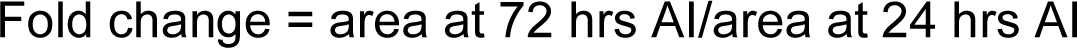

To count the epidermal cells, we counted 6 apodemes on either side of the hemisegment, centered around the neuron’s cell body, then drew “lines” between the two top apodemes and the two bottom apodemes. We counted all epidermal cells within the area created.

Fold change in F (fluorescence) was calculated using the following equation:

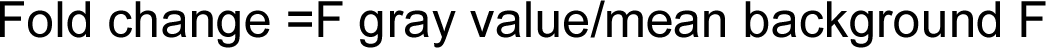

ΔF/F_background_ was calculated using the following equation:

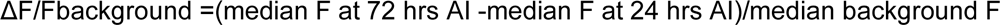

Sample sizes for all figures are as follows. Fig. 1E uninjured control n = 2, pinch n = 2; Fig. 1-1A 2p n = 3, pinch n = 3; Fig. 2B n = 10; Fig. 2C n = 11; Fig. 2D uninjured control n = 10, pinch n = 11; Fig. 2F n = 14; Fig. 2G n = 16; Fig. 2H uninjured control n = 14, pinch n = 16; Fig. 3C pinch n = 11, 2p n = 13; Fig. 3D pinch n = 11, 2p n = 13; Fig. 3G n = 9; Fig. 3H n = 9; Fig. 3-1B n = 12; Fig. 3-1C n = 13;Fig. 3-1D uninjured control n = 12, 2p n =13; Fig. 4B; Fig. 4C 2p n = 6, pinch n = 4; Fig. 4D uninjured control at 24 hrs AI n = 5, uninjured control at 72 hrs AI n = 3, pinch at 24 hrs AI n = 7, pinch at 72 hrs AI n = 4; Fig. 4E 2p n = 4, pinch n = 4; Fig. 4F uninjured control n = 5, 2p n = 4, pinch n = 4; Fig. 4G 2p n = 3, pinch n = 2; Fig. 4H 2p n = 5, pinch n = 7; Fig. 4-1A n = 7; Fig. 4-1B n = 3; Fig. 4-1C n = 8; Fig. 4-2A uninjured control at 24 hrs AI n = 8, uninjured control at 72 hrs AI n = 6; Fig. 4-2C pinch at 24 hrs AI n = 7, pinch at 72 hrs AI n = 4; Fig. 4-2E uninjured control at 24 hrs AI n = 5, pinch at 24 hrs AI n = 7, uninjured control at 72 hrs AI n = 3, pinch at 72 hrs AI n = 4; Fig. 4-2F 2p at 24 hrs AI n = 6, 2p at 72 hrs AI n = 6; Fig. 4-2G uninjured control at 24 hrs AI n = 3, 2p at 24 hrs AI n = 6, uninjured control at 72 hrs AI n = 3, 2p at 72 hrs AI n = 6; Fig. 4-3A n = 5; Fig. 4-3B n = 5; Fig. 4-3C n = 4; Fig. 4-3D n = 4; Fig. 4-3E n = 4; Fig. 4-3F n = 4; Fig. 4-3G n = 1; Fig. 4-4A n = 5; Fig. 4-4B n = 3; Fig. 4-4C n = 2; Fig. 4-5A n = 11; Fig. 4-5B n = 7; Fig. 4-5C n = 5.

**Figure 4:**
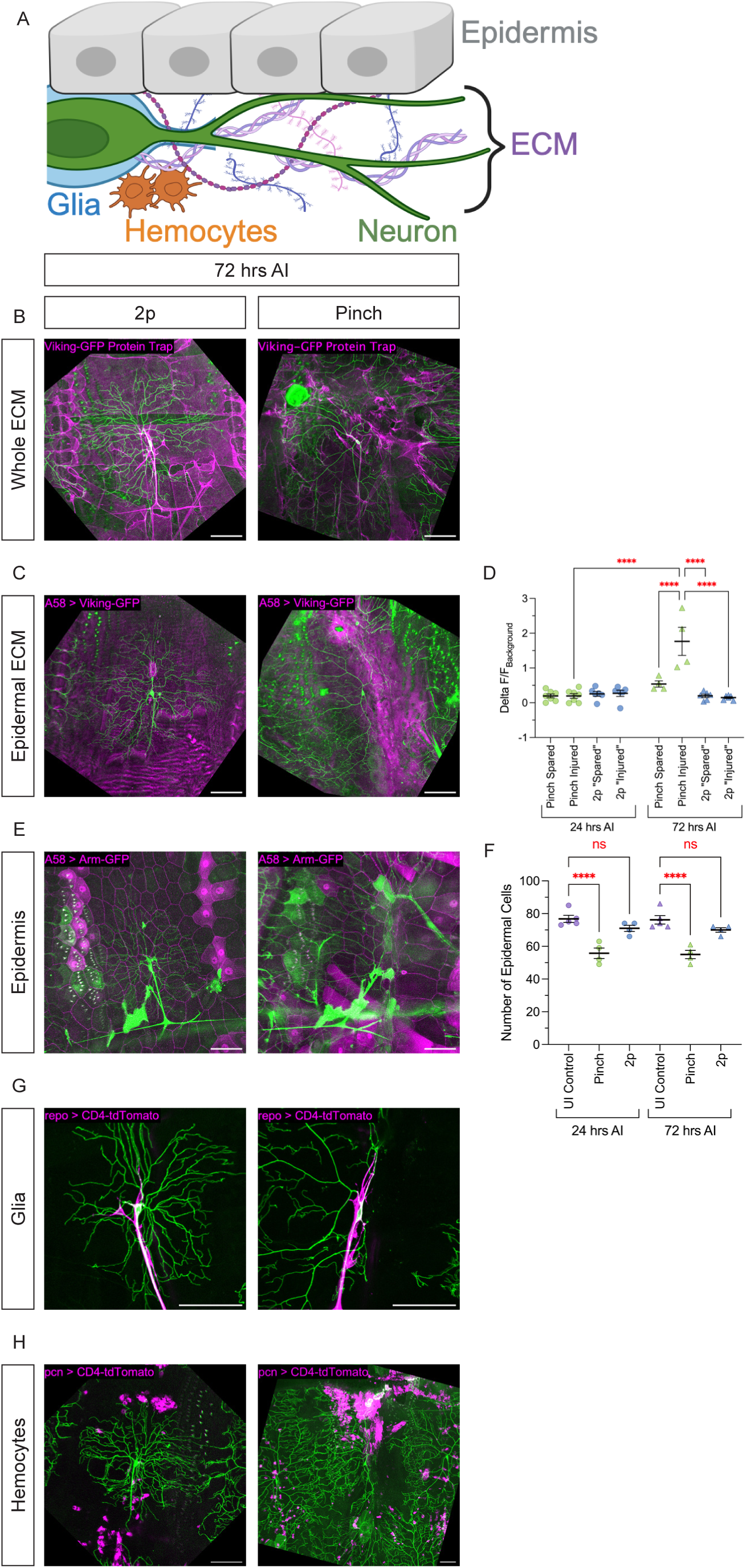
Pinch Injury Damages ECM, Epidermal Cells, and Recruits Hemocytes, but Does Not Damage Glia **(A)** Schematic of a class IV ddaC neuron (green) with the other cell types and tissues it interacts with: epidermis (gray), glia (blue), ECM (purple), and hemocytes (orange). **(B, C, E, G, H)** ddaC neurons (green) with other tissues in magenta following 2p-full bald (left) or pinch (right) injuries at 72 hrs AI. B: whole ECM protein trap (vkg-GFP); see Fig 4-1 for individual channels. C: A58-driven ECM (Gal4^A58^ > vkg-GFP); see Fig 4-2 for individual channels and quantification. E: epidermal tight junction marker (Gal4^A58^ > Armadillo-GFP); see Fig 4-3 for individual channels and counts of epidermal cells. G: glial marker (Gal4^repo^ > CD4-tdTomato); see Fig 4-4 for individual channels. H: hemocyte marker (Gal4^pcn^ > CD4-tdTomato); see Fig 4-5 for individual channels. Scale bar 100 µm. **(D)** ΔF/F_background_ of median fluorescent values of Gal4^A58^ > vkg-GFP following pinch (green) or 2p-full bald (blue) injuries at 24 and 72 hrs AI. Fluorescent values of pinched neurons are separated by pinched and spared sides. Fluorescent values of 2p-full bald neurons are arbitrarily separated into “injured” (right) and “spared” (left) sides. **(F)** Number of epidermal cells in uninjured (purple), pinched (green), or 2p-full bald (blue) neurons at 24 and 72 hrs AI.

All sample sizes for paired graphs of 24 to 72 hrs have the same n and are analyzing the same neurons at both time points, except for the datasets collected for the A58-driven ECM (a few of which are missing the 72 hr after injury time point). Nearly all injured neurons have a within-animal uninjured control neuron.

All images displayed are representative of the phenotype. Representative images were selected for their clarity.

### Some schematics were created with BioRender

Statistical analysis was performed using Graphpad Prism software (version 10.3.1). A two-tailed Welch’s student’s t-test was used for all analysis comparing two groups, using a paired student’s t-test for all analysis comparing the same neuron at two time points, using a one-way ANOVA with Holm-Šidák’s multiple comparisons correction for analysis comparing more than two groups, and using a two-way ANOVA with Holm-Šidák’s multiple comparisons correction for analysis with more than two variables. For all statistical tests, *p<0.05, **p<0.01, ***p<0.001, ****p<0.0001, and ^ns^p>0.05.

Statistical tests for all figures are as follows. Fig. 1E: unpaired t test; Fig. 2B: Number of Branch Tips = paired t test, Total Branch Length = paired t test; Fig. 2C: Number of Branch Tips = paired t test, Total Branch Length = paired t test; Fig. 2D: unpaired t test with Welch’s correction; Fig. 2F: Number of Branch Tips = paired t test, Total Branch Length = paired t test; Fig. 2G: Number of Branch Tips = paired t test, Total Branch Length = paired t test; Fig. 2H: unpaired t test with Welch’s correction; Fig. 3C: Number of Branch Tips = Two-way ANOVA with Holm-Šidák’s multiple comparisons test, Total Branch Length = Two-way ANOVA with Holm-Šidák’s multiple comparisons test; Fig. 3D: Unpaired t test with Welch’s correction; Fig. 3G: Number of Branch Tips 24 hrs AI = unpaired t test, Number of Branch Tips 72 hrs AI = unpaired t test, Total Branch Length 24 hrs AI = unpaired t test, Total Branch Length 72 hrs AI = unpaired t test; Fig. 3H: 24 hrs AI = unpaired t test, 72 hrs AI = unpaired t test; Fig. 4D: Two-way ANOVA with Holm-Šidák’s multiple comparisons test; Fig. 4E: Two-way ANOVA with Holm-Šidák’s multiple comparisons test; Fig. 1-1A: unpaired t test; Fig. 3-1B: Number of Branch Tips = paired t test, Total Branch Length = paired t test; Fig. 3-1C: Number of Branch Tips = paired t test, Total Branch Length = paired t test; Fig. 3-1D: unpaired t test with Welch’s correction; Fig. 4-2E: Two-way ANOVA with Holm-Šidák’s multiple comparisons test; Fig. 4-2G: Two-way RM ANOVA with Holm-Šidák’s multiple comparisons test; Fig. 4-3B: paired t test; Fig. 4 - 3D: paired t test. Fig. 4-3E: paired t test.

## Key Resources Table

| Reagent or Resource | Source | Identifier |
| --- | --- | --- |
| **Chemicals** |  |  |
| DuPont Molykote® High Vacuum Grease | Electron Microscopy Sciences | Cat. # 60705 |
| Glycerol ReagentPlus®, ≥99.0% (GC) | Sigma Aldrich | CAS # 56-81-5 |
| Isoflurane | Piramal, Midwest Vet Supply | Item# 193.33165.3 |
| GeneMate LE Quick Dissolve Agarose | VWR | Cat. # E3119-500 |
| **Experimental Models: Organisms/strains** |  |  |
| *Drosophila*: Canton-S | Lab Stock | RRID: FlyBase_FBsn0000274 |
| *Drosophila*: WeeP304 (tau-GFP) | Melissa Rolls, Michelle Stone et al. | RRID: FlyBase_FBal0249389 |
| *Drosophila*: w[1118]; P{w[+mC]=ppk-CD4-tdGFP}1b | BDSC | RRID:BDSC_35842,<br>FlyBase_FBrf0214034 |
| *Drosophila*: ; ppk-Gal4<sup>^</sup>VK34 / CyOw ; ppk-CD4-tdGFP / Tm6B | Lab Stock |  |
| *Drosophila*: w[1118]; P{y[+t7.7] w[+mC]=UAS-CaMPARI2.L398T}su(Hw)attP5 | BDSC | RRID:BDSC_78319,<br>FlyBase_FBTi0199488 |
| *Drosophila*: ; + ; Gal4<sup>^</sup>2-21, UAS-CD4-tdTomato / Tm6B | Lab Stock |  |
| *Drosophila*: w[1118]; P{w[+mC]=ppk-CD4-tdTom}4a | BDSC | RRID:BDSC_35844,<br>FlyBase_FBrf0214034 |
| *Drosophila*: ;; A58-Gal4 | Lab Stock | FlyBase_FBaI0181674 |
| *Drosophila*: y[1] w[*]; P{w[+mC]=PTT-un}vkg[G00454] | BDSC | RRID:BDSC_98343 |
| *Drosophila*: + ; sp / CyO ; repo-Gal4, UAS-mRFP / TM6B | Lab Stock |  |
| *Drosophila*: w[*]; M{w[+mC]=UASp-arm.mGFP6}ZH-86Fb/TM6B, Tb[1] | BDSC | RRID:BDSC_58725,<br>FlyBase_FBrf0227192 |
| *Drosophila*: ;; UAS-Viking-GFP / Tm3, Sb | Stéphane Noselli | FlyBase_FBtp0125825 |
| *Drosophila*: w[1118]; P{w[+mC]=Pxn-GAL4}3 | BDSC | RRID:BDSC_600223 |
| *Drosophila*: y[1] w[*]; P{w[+mC]=UAS-CD4-tdTom}7M1 | BDSC | RRID:BDSC_35841,<br>FlyBase_FBrf0214034 |
| **Software and Algorithms** |  |  |
| ImageJ (FIJI) | Wayne Rasband | RRID:SCR_002285<br>https://fiji.sc |
| Adobe InDesign | Adobe | RRID:SCR_021799<br>https://www.adobe.com/products/indesign.html |
| GraphPad Prism | GraphPad | RRID:SCR_002798<br>https://www.graphpad.com |
| Zen Microscopy Software | Zeiss | RRID:SCR_013672<br>https://www.zeiss.com/microscopy/en/products/software/zeiss-zen.html |
| Microsoft Excel | Microsoft | RRID:SCR_016137<br>https://www.microsoft.com |
| **Other** |  |  |
| LSM 700 Fluorescent Confocal Microscope | Zeiss | RRID:SCR_017377 |
| LSM 980 Fluorescent Confocal Microscope with Airyscan 2 | Zeiss | RRID:SCR_025048 |
| MAI TAI™ 2-photon laser | Spectra-Physics | Model EHP 1040S |
| Dissecting Microscopes (Zeiss Stemi 2000, Nikon SMZ-10A, Olympus SZ60) | Zeiss, Nikon, Olympus | Models Stemi 2000; SMZ-10A; SZ60 |
| Fiber-Lite Mi-150 Illuminator Series, 150w Halogen light source (MI150 Mi 150) | Dolan-Jenner Industries | Cat. # 660000391010 |
| Eppendorf™ Micropipettes | Thermo Fisher Scientific | SKU P-10MLR |
| Epredia™ Richard-Allan Scientific™ Cover Glass 22mm x 22mm | Thermo Fisher Scientific | Cat. # 22-050-235 |
| Fisherbrand™ Premium Plain Glass Microscope Slides | Thermo Fisher Scientific | Cat. # 12-544-1 |
| LEICA MZ FLIII Fluorescent sorting microscope | LEICA | Model MZ FLIII |
| Super High Pressure Mercury Lamp Power Supply | Nikon | Model HB-10101AF |
| Fisherbrand™ Round Bottom Disposable Borosilicate Glass Tubes with Plain End | Thermo Fisher Scientific | Cat. # 14-961-27 |
| Parafilm® M Sealing Film | Sigma Aldrich | Cat. # HS234526B-1EA |
| Syringe, 10 ml, 5ml | Thermo Fisher Scientific | Cat. # 14-955-458; Cat# 14-955-459 |
| *Drosophila* food | UCI Fly Food Kitchen | N/A |
| Yeast, Active, Dry | Fisher Science Education | Cat. # S25632 |
| Welch's 100% Grape Juice | Welch's | WEL35400 |
| Drosophila Incubator | Percival | https://www.percival-scientific.com/product-category/drosophila/ |
| Handheld UV lamp | Maxxeon | MXN02003 |

## Data Availability

The datasets generated and/or analyzed during the current study are available from the lead contact upon reasonable request. This paper does not report original code. Any additional information required to reanalyze the data reported in this paper is available from the lead contact upon request.

## Results

### Pinch injury damages da PNS neurons in *Drosophila melanogaster* larvae

To develop an injury model that better mimics widespread tissue damage, we adapted a pinch injury method established by Burra, et al (Burra et al., 2013). We modified the injury method such that cuticular puncture did not occur (Figure 1A). To do this, we blunted a pair of #5 forceps and pinched the tissue laterally such that the larva’s cuticle and epithelia bulged out between the tines of the forceps, which we termed a “larval muffin top” (Figure 1A and 1B). The pinch injury was held for 10 seconds and typically lifted the larvae up into the air (Figure 1B and Video 1). Compared to the traditional dendrite severing by a 2-photon (2p) laser, our new pinch injury method is significantly faster to perform (Figure 1-1; 17.2 minutes per injury for 2p versus 4.2 minutes per injury for pinch).

Following pinch injury, we first inquired about the extent of damage the class IV da neuron, ddaC, sustained. We visualized neurons using the cell-type specific expression of a membrane-tagged GFP and imaged them at 24 hours after pinch. We found that abdominal segment ddaC neuron A4 or A5 was injured by our approach: one lateral half of the dendritic arbor was damaged, and the other half was left intact (Figure 1C).

Next, we set out to determine whether pinch injury led to calcium influx, the first step of dendrite injury detection in da neurons (Duarte et al., 2024; Hertzler et al., 2024). We used CaMPARI (Calcium-Modulated Photoactivatable Ratiometric Indicator), a photoactivatable calcium indicator that photoconverts from green to red only in the simultaneous presence of calcium ions and UV light (Fosque et al., 2015). Immediately following pinch injury, larvae expressing CaMPARI in ddaC neurons were exposed to 20 seconds of UV light. Red converted CaMPARI fluorescence was detected in the soma and proximal primary dendrites in pinched ddaC neurons, but not in uninjured (UI) control neurons (Figure 1D). When we quantified this effect, pinched ddaC neurons had a significantly higher red fluorescence intensity of converted CaMPARI compared to uninjured controls (Figure 1E). These results indicate that there is calcium influx into ddaC neurons following pinch injury, which is a critical step in dendrite injury detection and is similar to what has been observed immediately following 2p injury (Duarte et al., 2024).

Together, these results demonstrate that pinching injury can damage dendrites, in a highly reproducible and rapid manner.

### ddaC and ddaE neurons (re)grow their branches following pinch injury

To assess how da neuronal dendrites respond to pinch injury, we first decided to evaluate the dendritic arbors of the class IV ddaC neurons. The dendritic arbors of class IV ddaC neurons are large, filling their receptive field by 48 hours after egg lay (AEL) and are very dynamic (Grueber et al., 2002; Parrish et al., 2009). Further, they retain a robust capacity for dendrite regeneration (Song et al., 2012; Stone et al., 2014; Thompson-Peer et al., 2016; Hertzler et al., 2023; Duarte et al., 2024; Prange et al., 2024; Hwu et al., 2025). Pinch injury damages half of the dendrite arbor and leaves the other half uninjured (Figure 2A). Following pinch injury, the spared half of the arbor becomes visibly bushier as it gains branches over time (Figure 2A). Specifically, ddaC neurons increase in number of branches and total dendrite length from 24 to 72 hrs after injury (AI) (Figures 2A and 2C). The growth on the spared half of the arbor is roughly similar to uninjured ddaC neurons (Figures 2A and 2B), which continue to gain branches over time as the larvae grow and develop (Parrish et al., 2009).

In order to determine if the branches gained on the spared half of the pinched neuron was truly regeneration or simply the branch growth of development, we took two approaches. First, we looked at class I ddaE neurons, which are neurons that establish their arbors by 24 hrs AEL and are very static during later development (Sugimura et al., 2003). In the absence of injury, they do not change in branch number over time, though they do increase in branch length as the larvae grow (Figures 2E and 2F). Thus, any significant increase in branch number following pinch injury would be indicative of regeneration. We found that pinched class I ddaE neurons do increase in branch number and total dendrite length from 24 to 72 hrs after pinch, compared to the stable uninjured neurons that do not increase in branch number during this time (Figures 2E and 2G). As a second approach, we evaluated the growth rate of each neuron subtype, both class IV ddaC and class I ddaE neurons. We found that both have greater fold changes in branch number (number of branches at 72 hrs AI divided by number of branches at 24 hrs AI) following pinch injury compared to their uninjured controls (Figures 2D and 2H).

This result coupled with the regenerative response of class I ddaE neurons following pinch injury is indicative that both class IV ddaC and class I ddaE neurons exhibit regeneration of their dendrites following pinch injury and not just continued developmental growth.

### ddaC neurons preferentially regenerate into undamaged tissue

Next, we wanted to compare regeneration following pinch injury versus 2p injury, as precise dendrite severing by a 2p laser is the “gold standard” for precise injury in the field (Hertzler and Rolls, 2024). To compare the two injury methods, we mimicked the amount of dendrite removal obtained with a pinch injury by using a 2p laser to cut off half of the class IV dendritic arbor (Figure 3A and 3B and Figure 3-1A). We called this injury a ‘half-2p bald’ as half of the neuron was left “bald” by removal of half of its arbor. Class IV ddaC neurons regenerate following this half-2p bald laser injury. They increase their number of branches and total branch length, and they have a significantly higher fold change in branch number compared to their UI controls (Figure 3B, Figures 3-1B, 3-1C, and 3-1D).

When comparing the total number of branches and total dendrite length between pinched and half-2p balded class IV ddaC neurons, there are no significant differences at 24 or 72 hrs AI in the number or total length of dendrites (Figure 3C). We noticed, however, that pinch injury resulted in less dendrites innervating the empty territory on the injured side relative to half-2p balded neurons (black boxes in Figure 3B). Since this phenotype was not captured by calculating the total number of branches or total dendrite length (Figure 3C), we decided to calculate the fold change in area coverage of only the injured half of the arbor. Neurons injured by half-2p bald had a greater fold change in area coverage, reflecting our observation of an increased capacity for innervating empty territory following injury (Figure 3D).

Given this difference in regenerative capacity, we next wanted to challenge individual neurons to regenerate their dendrites following both pinch and 2p injury. Class IV ddaC neurons were first pinched and 1-2 hours later, the remaining arbor, the other half, was cut off with the 2p laser. This specific “half-pinch + half-2p” injury paradigm removed all branches from the class IV ddaC neuron and forced new dendrites to choose to regrow on one side or the other (Figures 3E and F). For all quantifications of this injury paradigm, the axon was used as a guideline to separate the two halves of the arbor. Following this half-pinch + half-2p injury, there was no statistical difference between the two sides in branch number, total dendrite length, or area coverage at 24 hrs AI (Figures 3G and 3H). However, at 72 hrs AI, the half of the arbor responding to 2p injury regenerated to significantly more, with respect to branch number, total dendrite length, and area coverage, than the half of the arbor responding to pinch injury (Figures 3G and 3H).

Taking the results of these two experiments together, we can conclude that ddaC neurons preferentially regenerate on the side injured by 2p injury. Since challenging individual neurons to regenerate following both injury paradigms led to differential regenerative outcomes, we hypothesized that extrinsic mechanisms may be causing poor re-innervation following pinch injury, either by active inhibition or the absence of support.

### Pinch injury damages other cell types and surrounding tissues

Following the differential regeneration of dendrites observed from the half-pinch + half-2p experiments, we wanted to evaluate how the extracellular environment might be affecting regeneration following both injury paradigms. There are a number of cell types and tissues that interact with the da neurons (Figure 4A). The da neuronal dendrites are suspended in an extracellular matrix (ECM) and innervate the larval epidermis. Glia wrap the axon, cell body, and proximal dendrites of the da neurons, and resident immune cells, hemocytes, are present. First, we examined the ECM.

Throughout development, the larval da neurons adhere to components of the ECM through adhesion molecules like integrins (Kim et al., 2012). These interactions help constrain dendrites to a 2D space: within the ECM and between the musculature and epidermis (Grueber et al., 2002; Yasunaga et al., 2010; Han et al., 2012; Kim et al., 2012). Major components of the ECM include laminins and collagen IV, which are primarily produced by the fat body and hemocytes, but also by the larval epidermis (Rodriguez et al., 1996; Han et al., 2012; Isabella and Horne-Badovinac, 2015; Ramos-Lewis et al., 2018).

Thus, to visualize the ECM, we chose to look at viking (vkg), which is homologous to the mammalian Collagen IV (Yasothornsrikul et al., 1997). We started off using a protein trap fly line where the endogenous vkg gene is tagged with GFP. For uninjured neurons, vkg-GFP predominantly wraps the muscle and muscle attachment sites; it also wraps the axon, cell body, and proximal dendrites of the ddaC class IV neurons (Figure 4-1A). Following 2p injury, vkg-GFP is undamaged (Figure 4B and Figure 4-1C). Following pinch injury, vkg-GFP is severely damaged (Figure 4B and Figure 4-1B). The vkg-GFP wrapping the muscle and muscle attachment sides are significantly damaged, and there is no longer a consistent vkg-GFP signal across the hemisegment. Vkg-GFP continues to wrap the axon, cell body, and proximal dendrites of the spared half of the pinched arbor. However, there is a significant accumulation of vkg-GFP signal on the pinched half of the arbor that aligns with the pinched edge of the injured side of the arbor. This accumulation of vkg-GFP is reminiscent of a glial or fibrotic scar, which can create an inhibitory, non-permissive environment for neurite regrowth (Liesi and Kauppila, 2002; Klapka and Müller, 2006; Tran et al., 2022).

Since the whole-body vkg-GFP has a significant amount of signal, mostly surrounding the muscle, it is difficult to see the ECM directly interacting with the dendrites. Thus, we sought to visualize vkg specifically expressed by A58-Gal4, which drives expression in the larval epidermis and, in later third instar stages, the fat body (Ramos-Lewis et al., 2018). Uninjured hemisegments have a smooth, even vkg-GFP fluorescence (F) signal that mimics the “cobblestone” pattern of the epidermis (Figure 4-2A, quantified in Figure 4-2B). The A58-driven vkg-GFP appears unaltered by 2p injury (Figure 4C and Figure 4-2F, quantified in Figure 4-2G). Thus, the A58-driven vkg-GFP is uninjured following 2p injury. Following pinch injury, we found that A58-driven vkg-GFP is enriched on the pinched side of the arbor compared to the spared side or an uninjured neuron or a neuron injured by 2p (Figure 4C and Figure 4-2C). The enrichment of A58-driven vkg-GFP on the pinched half of the neuron increases from 24 to 72 hrs AI (Figure 4-2D and E). This is reminiscent of the enrichment of whole-body vkg-GFP observed in Figure 4B. As an additional note, we observed a slight increase in the A58-driven vkg-GFP fluorescence on the spared half at 72 hrs AI, most likely due to diffusion of vkg-GFP from the pinched side (Figure 4D and Figure 4-2E). Thus, both endogenous and over-expressed viking accumulates and persists at the site of injury, only after pinch injury.

Next, we wanted to look at the larval epidermis. The da neurons establish their arbors in a 2D space: within the ECM and between the musculature and epidermis (Grueber et al., 2002; Yasunaga et al., 2010; Han et al., 2012; Kim et al., 2012). The larval epidermis is a monolayer sheet of terminally differentiated epidermal cells, and plays a crucial role in establishing the dendritic arbor of the da neurons (Parrish et al., 2009; Burra et al., 2013; Jiang et al., 2014; Poe et al., 2017). HSPGs expressed by the larval epidermis are required for stabilization of high-order dendrites in ddaC class IV neurons (Poe et al., 2017). Additionally, portions of ddaC class IV dendrite branches are enclosed by epidermal cells via coracle to restrict dendrite outgrowth and mediate the occupancy of multiple classes of da neurons in a 2D space (Tenenbaum et al., 2017). To visualize the epidermis, we used a GFP-tagged version of Armadillo, the fly homolog of beta-catenin and an adherens-junction marker (Cox et al., 1996). For uninjured neurons, the epidermis is a single sheet of polygonal epidermal cells (Figure 4-3A), and we found no difference in the number of epidermal cells from 24 to 72 hrs AI (Figure 4-3B). Following 2p injury, the epidermis appears the same as uninjured hemisegments (Figure 4E and Figure 4 - 3E), and we also found no difference in the number of epidermal cells from 24 to 72 hrs AI (Figure 4-3F). Further, there is no statistical difference between the number of epidermal cells in uninjured or 2p-injured neurons at 24 or 72 hrs AI, further supporting that there are no differences in the epidermis caused by 2p injury (Figure 4F). However, following pinch injury, the epidermis is damaged (Figure 4E and Figure 4-3C). On the pinched side of the neuron, the epidermal cells are disfigured. Close to the focal point of injury, the epidermal cells are small and crushed. Next to the muscle attachment sites (apodemes), the epidermal cells are elongated. On the spared side, the epidermal cells appear uninjured, similar to uninjured and 2p-injured hemisegments. There is no statistical difference between the number of epidermal cells at 24 or 72 hrs AI following pinch injury (Figure 4-3D). However, there are significantly fewer epidermal cells following pinch injury than there are for uninjured neurons, indicating that the epidermal cells were damaged and some died following pinch injury (Figure 4F). Damage to epidermal cells may contribute to why pinch-injured neurons exhibit poor regrowth into empty territory.

Next, we looked at the glia. Dendritic arborization neurons have three layers of glia that wrap the axon, the soma, and the proximal dendrites (Yadav et al., 2019). Glia are indispensable for establishing both the structure and function of ddaC class IV neurons. Uninjured neurons have glia that wrap the axon and somatodendritic compartments - specifically, the proximal region of the primary dendrites (Figure 4-4A) (Yadav et al., 2019). Following 2p injury, the glia are capable of wrapping the axon and soma (Thompson-Peer et al., 2016), but the glia have trouble wrapping the proximal segments of the newly regenerated primary dendrites (Figure 4G and Figure 4-4C). Following pinch injury, the glia show no signs of obvious injury (Figure 4G and Figure 4-4B). On the spared side of the arbor, the glia appear normal; on the pinched side of the arbor, the primary dendrites and glial wrapping are absent. The glia can, however, still wrap the axon and soma following pinch injury. Thus, glia are mostly unaffected following either injury paradigm.

Lastly, we looked at the immune response following both injury paradigms. There are three types of immune cells (hemocytes) in *Drosophila*: plasmatocytes, crystal cells, and lamellocytes (Rizki, 1957; Rizki and Rizki, 1983, 1992; Tepass et al., 1994; Lebestky et al., 2000). Approximately 95% of all circulating hemocytes are plasmatocytes, and the remaining 5% are crystal cells. Lamellocytes are only stimulated by parasitism (Rizki and Rizki, 1992; Meister and Lagueux, 2003). While most hemocytes migrate through the hemolymph and have no specific localizations, in abdominal segments 1-7 (in which these injury experiments were performed), there are some specific groups of localized hemocytes in the epidermal-muscular pocket (Makhijani et al., 2011). Proper localization and survival of these resident hemocytes is supported by the da neurons (Makhijani et al., 2011).

These resident hemocytes include the dorsal vessel cluster (near the top of the ddaC class IV dendritic arbor), the dorsal stripe (near the soma), and the lateral patch (which extends from the bottom of the ddaC class IV dendritic arbor into the neighboring v’ada class IV arbor) (Makhijani et al., 2011). These three patches of hemocytes are visible in uninjured hemisegments (Figure 4-5A). Following 2p injury, there is no major accumulation of hemocytes near the regenerating arbor, and the three patches of hemocytes appear normal (Figure 4H and Figure 4-5C). Following pinch injury, however, there is major accumulation of hemocytes on the pinched side of the ddaC class IV arbor (Figure 4H and Figure 4-5B). There are no longer three discrete patches of hemocytes; instead, there is a large influx of hemocytes near the focal point of injury. This influx of hemocytes at the site of injury points towards a large immune reaction following the wide-spread tissue damage caused by pinch injury, representing another possible reason for poor dendrite regrowth into empty territory.

## Discussion

Here, we have established a pinch injury method to injure dendrites that causes holistic tissue damage, effectively damaging the neurons, ECM, and epidermis, while leaving the glia largely intact. This pinch injury method elicits robust immune infiltration to the injured half of the dendritic arbor. Importantly, the surrounding tissue damage restricts invasive dendrite regeneration into the empty territory, instead driving compensatory regeneration on the spared side of the arbor. This contrasts sharply with the invasive dendritic growth observed following 2p laser injury, where the epidermis and ECM remain undamaged. The discrete regenerative response to these two injury paradigms was further demonstrated in our half-pinch/half-2p injury paradigm: the pinched half of the class IV neuron exhibited reduced branch number, reduced total branch length, and reduced area coverage compared to the 2p-injured half of the dendritic arbor. These findings highlight that an intact and healthy extracellular environment is critical for dendrite regeneration. Given that the pinch injury damages not only neurons but also surrounding tissue, analogous to what occurs in traumatic brain injury (TBI), blunt force trauma, sports-related injuries, and blast injuries, this model provides a physiologically relevant platform to study neuronal regeneration (Hicks et al., 2010; Mustafa and Alshboul, 2013; George and Geller, 2018). In these real-world injuries, damage extends beyond neurons to include the ECM, supportive glial cells, and endothelial cells comprising the blood-brain barrier, making the pinch injury a more accurate model of acute neurotrauma than laser ablation alone (Gaudet et al., 2011; George and Geller, 2018).

The pinch injury method we describe here complements existing injury paradigms used to study neuronal regeneration. While laser ablation (either 2p or pulsed UV laser) has been instrumental in advancing our understanding of dendrite regeneration due to its precision and reproducibility (Duarte et al., 2024; Hertzler and Rolls, 2024; Prange et al., 2024; Hwu et al., 2025), it is fundamentally different from physiological injury in that it leaves surrounding tissue intact. Mechanical injury models, such as axon crush injuries commonly used to study peripheral nerve regeneration, cause widespread tissue damage similar to our pinch injury. Axon crush paradigms in *Drosophila* and vertebrate models have been widely used to study axon regeneration and have revealed that both the neuron and its local environment must be considered when evaluating regenerative capacity (Menorca et al., 2013; Rooney and Freeman, 2014; Akram et al., 2022; Bhattacharya, 2023; Pluta et al., 2025; Waller et al., 2025). Similarly, peripheral nerve injuries in mice, including sciatic nerve crush and pinch injuries, cause damage to neurons, Schwann cells, blood vessels, and ECM, triggering inflammatory responses that influence regeneration outcomes (Gaudet et al., 2011). Our dendrite pinch injury model parallels these axon injury paradigms but allows for spatial compartmentalization of damage to part of a single neuron. This unique feature (injuring only half of the dendritic arbor) enables direct within-neuron comparisons of regeneration in damaged versus undamaged tissue environments, providing insights that are harder to glean from whole-nerve injury models.

The accumulation of the *Drosophila* homolog of collagen IV, viking (vkg), and damage to the ECM that we observed following pinch injury may represent a scar-like structure that impedes dendrite regeneration. This vkg accumulation and scar-like structure is similar to the collagenous scar that forms in the ECM following dorsal pinch injury in *Drosophila* larvae (Ramos-Lewis et al., 2018). Further, in mammalian spinal cord injury (SCI) and TBI, glial scars form at the lesion site, characterized by reactive astrocytes, the deposition of chondroitin sulfate proteoglycans (CSPGs), and ECM remodeling (Siddiqui et al., 2022; Tran et al., 2022). Similarly, fibrotic scars composed of collagen and fibronectin create physical and chemical barriers to axonal regeneration following nerve injury (Ayazi et al., 2022). The vkg accumulation we observe in the pinched tissue shares features with these injury-induced scars, raising the possibility that it serves as a barrier to dendrite regrowth. However, the mechanism by which this damaged ECM might restrict dendrite regeneration remains unclear. Injured dendrites might be actively repelled by molecular cues within the damaged tissue, analogous to how CSPGs inhibit axon growth in the glial scar through receptor-mediated signaling (Ohtake and Li, 2015). Alternatively, the damaged epidermis and disrupted ECM architecture may simply fail to provide the adhesive substrate or trophic support necessary for dendrite attachment and growth. Distinguishing between active repulsion and failure of permissive attachment is critical for understanding whether dendrites avoid the injured territory or are unable to invade it. Our half-pinch/half-2p injury paradigm, which reveals preferential growth on the 2p-injured side, suggests that the lack of tissue damage may be sufficient to promote regeneration, favoring a model in which the damaged tissue fails to support dendrite growth rather than actively repelling it.

The tissue damage and inflammatory response following pinch injury are not static but rather evolve over time, potentially creating an increasingly restrictive environment for dendrite regeneration. We observe that vkg accumulation in the damaged ECM increases progressively following injury. This temporal pattern suggests that the barriers to regeneration may intensify as time progresses, rather than resolving to create a more permissive substrate for growth. The dynamic remodeling of the ECM following injury may be mediated in part by matrix metalloproteinases (MMPs), which degrade ECM components and play complex, time-dependent roles in regeneration (Yong, 2005; Phillips et al., 2014). In mammalian CNS injury, MMPs are rapidly upregulated in the acute phase and contribute to both beneficial effects, such as clearing debris and allowing cell migration, and detrimental effects, including disruption of the blood-brain barrier and exacerbation of inflammation (Andries et al., 2017). As such, the accumulation of vkg that we observe may reflect an active remodeling and reorganization of the ECM by MMPs, which in turn, could progressively limit the ability of the ECM to form a permissive substrate for dendrite growth (DeVault et al., 2018; Furusawa and Emoto, 2021).

While we have identified damage to the epidermis and ECM as key features of the pinch injury, the full extent of tissue damage and the cellular sources of ECM remodeling remain to be fully characterized. We used the Gal4^A58^ line to visualize vkg accumulation following injury, and while Gal4^A58^ is expressed in epidermal cells, it is also expressed in the fat body (Ramos-Lewis et al., 2018). Therefore, the vkg accumulation we observe may derive not only from epidermal cells but also from the fat body, which serves as a major source of ECM proteins and circulating factors in *Drosophila* (Rodriguez et al., 1996; Isabella and Horne-Badovinac, 2015; Ramos-Lewis et al., 2018). Distinguishing between these sources will be important for understanding the origin and regulation of the ECM response to injury. Moreover, vkg/collagen IV is just one of many ECM components, which we selected due to its high abundance and structural importance (Rozario and DeSimone, 2010). Other ECM components including laminins, proteoglycans, and other collagens, are likely also disrupted or dysregulated following pinch injury. The complete ECM composition in the injured territory, and how it differs from uninjured tissue, may provide insight into what makes the environment restrictive for dendrite growth. The pinch injury damages all cells within the compressed tissue, not only the sensory neuron and individual other cell types we imaged here. It is possible we are damaging or killing other da neurons and cell types that normally provide trophic support, guidance cues, or other pro-regenerative signals that facilitate dendritic outgrowth. Uncharacterized aspects of the injury underscore the complexity of the tissue damage and highlight the need for comprehensive characterization.

Lastly, an important consideration is how the combined damage to neurons and surrounding tissue would affect sensory function. Both class IV da neurons and epidermal cells are mechanosensitive and work cooperatively to detect mechanical stimuli in *Drosophila* larvae (Luedke et al., 2024; Yoshino et al., 2025). Damage as we observe following pinch injury could compromise mechanosensation through multiple mechanisms, including direct loss of epidermal mechanosensory function and disruption of neuron-epidermis communication (Yang and Chien, 2019; Luedke et al., 2024; Yoshino et al., 2025). This has important implications for understanding recovery from injury in clinical cases, where damage to neurons, glia, and other supporting cell types occurs simultaneously. Even if dendrites successfully grow new branches after pinch injury - as we observed through compensatory growth on the uninjured side of the arbor - full restoration of sensory function might require neuronal regrowth as well as repair of epidermal cells and the re-establishment of neuron-epidermis interactions.

### The authors would like to acknowledge the following funding sources

Research in this publication was supported by NIH Grant R00NS097627 and a UCI startup grant (to KTP). Research reported in this publication was also supported by the Department of Education’s Graduate Assistance in Areas of National Need (GAANN) Program P200A220015 and the Rose Hills Foundation Science & Engineering Fellowship. We would like to acknowledge Lauren Kim for her beautiful artistry while generating most of the schematics seen in this paper. We would also like to thank Patrick Hwu for his invaluable mentorship and scientific guidance throughout the project. We also thank Vinicius Duarte, Rosty Brichko, and Yuting Liu for their intellectual contributions to this work. The authors would also like to acknowledge the University of California Irvine’s Undergraduate Research Opportunities Program and Summer Undergraduate Research Program (UROP and SURP). Stocks obtained from the Bloomington Drosophila Stock Center (NIH P40OD018537) were used in this study. The authors would like to thank Stephane Noselli for graciously sending us a copy of the UAS-viking-GFP stock. We thank the UCI Optical Biology Core and Dr. Adeela Syed for extensive use of the microscopes in their facility. This study was made possible in part through access to the Optical Biology Core Facility of the Developmental Biology Center, a shared resource supported by the Cancer Center Support Grant (CA-62203) and NIH-S10OD032327-01. KTP is a fellow of the Hellman Foundation and the Rose Hills Foundation.

## Conflict of Interest

a. No; Authors report no conflict of interest Funding Sources:
b. NIH Grant R00NS097627
c. UCI startup grant (to KTP)
d. Department of Education’s Graduate Assistance in Areas of National Need (GAANN) Program P200A220015
e. Rose Hills Foundation Science & Engineering Fellowship
f. Bloomington Drosophila Stock Center (NIH P40OD018537)
g. Optical Biology Core Facility of the Developmental Biology Center, a shared resource supported by the Cancer Center Support Grant (CA-62203) and NIH-S10OD032327-01

**Video 1:** Video of Pinch Injury Method This video shows the positioning of the third-instar *Drosophila* larvae in preparation for pinching. It also shows the alignment of the two tines of the forceps with abdominal hemisegments 4 and 5, along with the slight “scooping” motion required to pinch the two hemisegments together.

---

**Figure 1-1:**
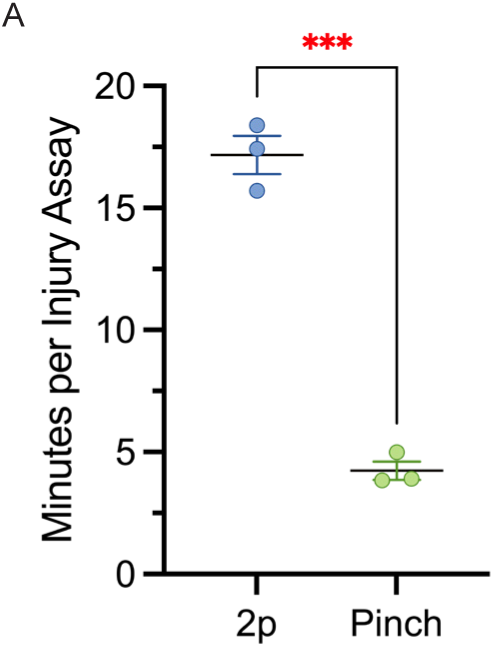
Pinch Injury Technique Is Faster to Perform Than 2p Dendrite Severing **(A)** Minutes required to perform injury to 1 neuron per animal for 2-photon (2p) injury assay (blue) and pinch injury assay (green).

**Figure 3-1:**
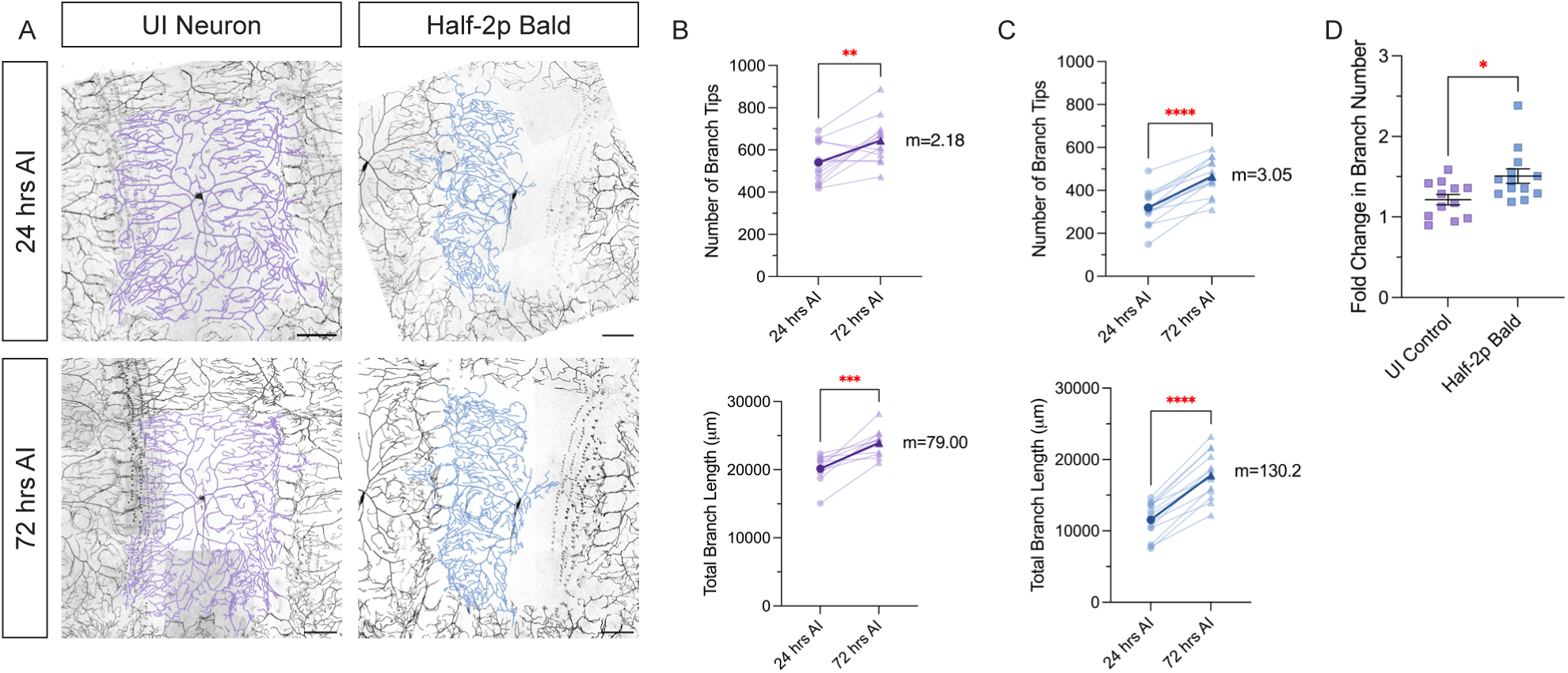
Half-2p Bald class IV ddaC Neurons Grow into Undamaged Tissue **(A)** Class IV ddaC uninjured (UI) within-animal control and half-2p injured neurons at 24 and 72 hrs AI. Scale bar 100 µm. **(B-C)** Number of branch tips (top) and total branch length (bottom) of uninjured (UI) within-animal control (B, purple) and half-2p bald (C, blue) neurons 24 to 72 hrs AI. **(D)** Fold change in branch number of uninjured (UI) within-animal control (purple) and half-2p bald (blue) class IV ddaC neurons.

**Figure 4-1:**
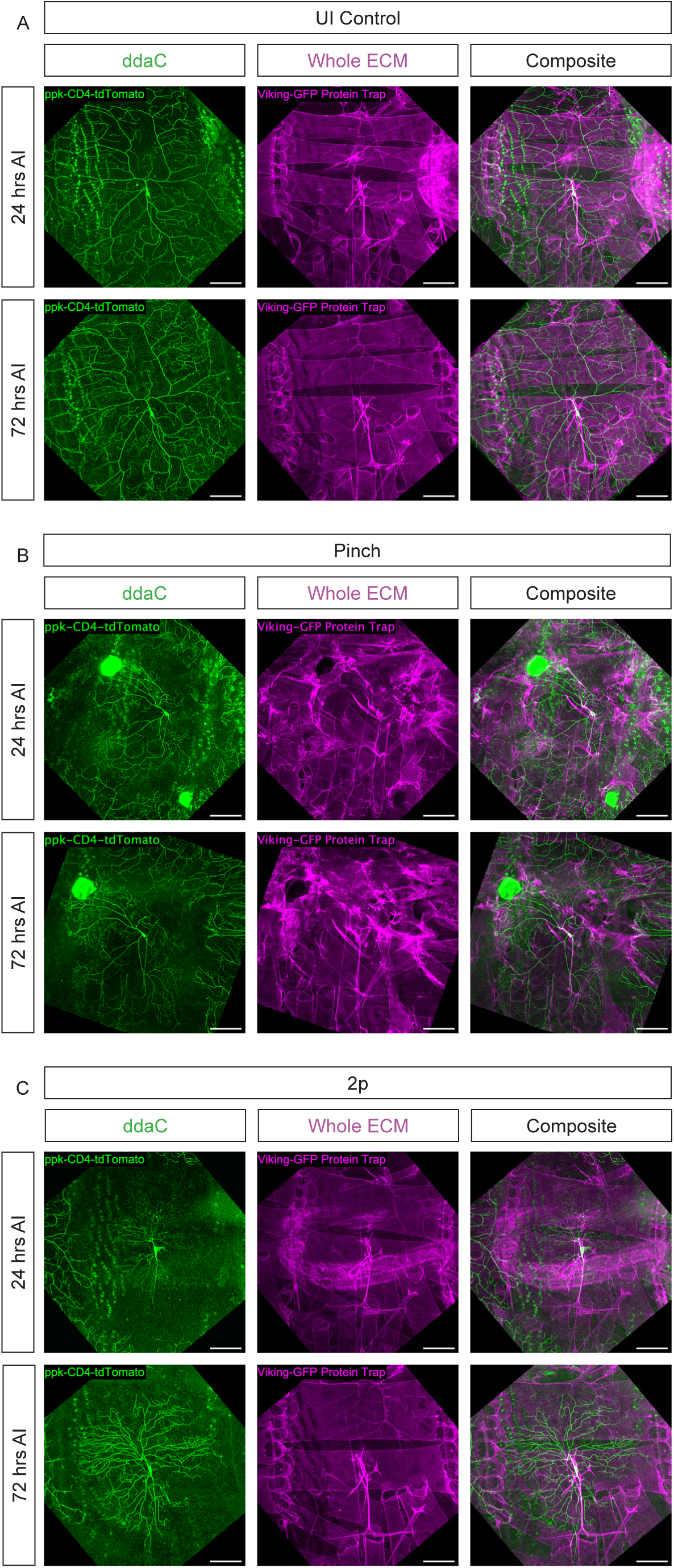
Pinch Injury, but Not 2p-Full Bald Injury, Damages Whole ECM viking/Collagen IV. (A-C) Uninjured (A), pinched (B), or 2p-full bald (C) class IV ddaC neurons with an ECM Protein Trap (vkg-GFP) at 24 and 72 hrs AI. Scale bar 100 µm. Composite images of pinched or 2p-full bald neurons at 72 hrs AI are also shown in Fig 4B.

**Figure 4-2:**
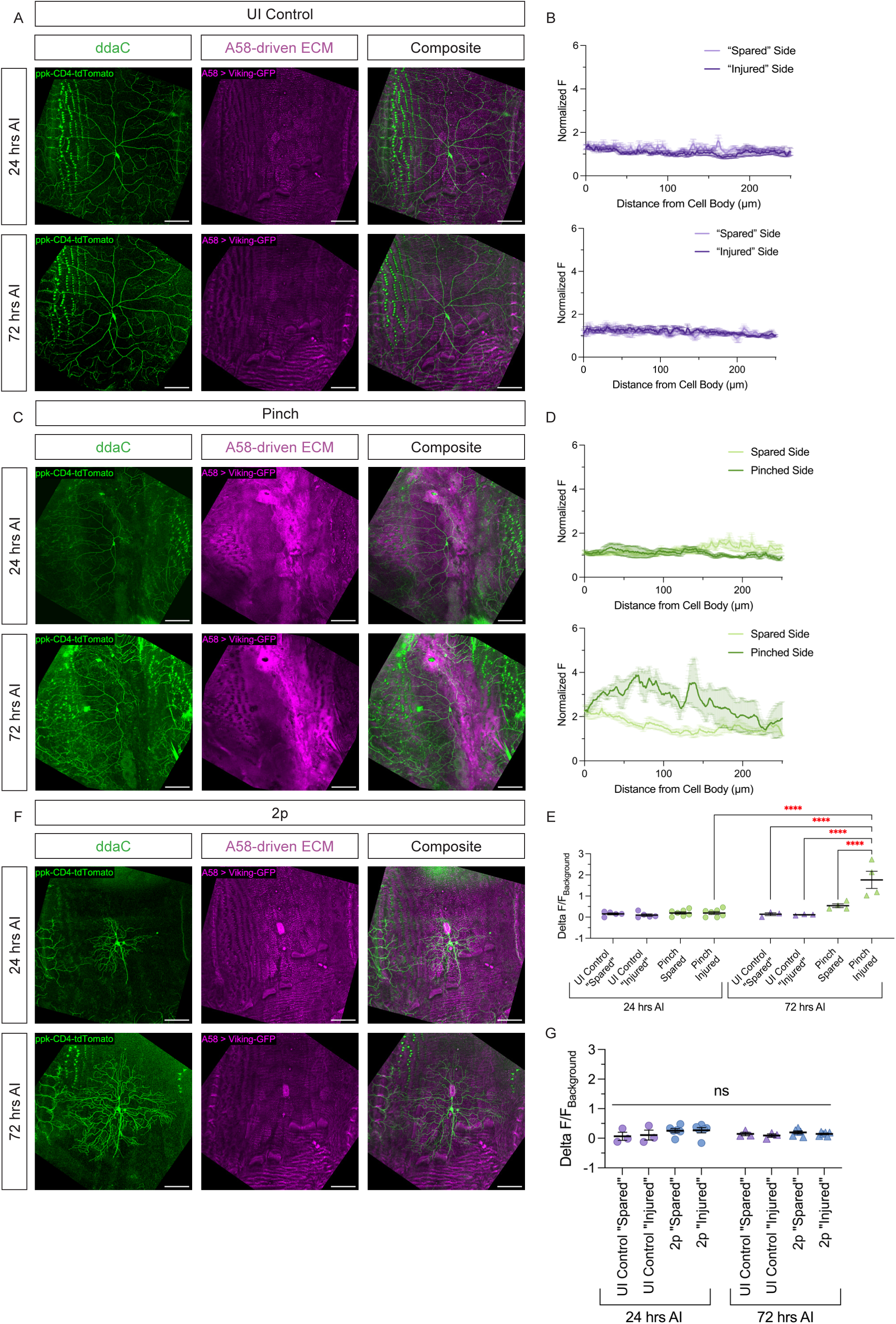
Pinch Injury, but Not 2p-Full Bald Injury, Damages A58-Driven ECM viking/Collagen IV. **(A, C, F)** Uninjured (A), pinched (C), or 2p-full bald (F) class IV ddaC neurons with A58-driven ECM marker (Gal4^A58^ > vkg-GFP) at 24 and 72 hrs AI. Composite images of pinched or 2p-full bald neurons at 72 hrs AI are also shown in Fig 4C. Scale bar 100 µm. **(B, D)** Normalized F (vkg-GFP fluorescence) over 250 µm starting from the cell body at 24 hrs AI (top) and 72 hrs AI (bottom), on the “spared” and “injured” sides of an uninjured neuron (B) or the spared and pinched sides of a pinched neuron (D). **(E, G)** ΔF/F_background_ of median fluorescence values of vkg-GFP at 24 and 72 hrs AI in uninjured (purple) or pinched (green) neurons (E), or uninjured (purple) and 2p-full bald (blue) neurons (G).

**Figure 4-3:**
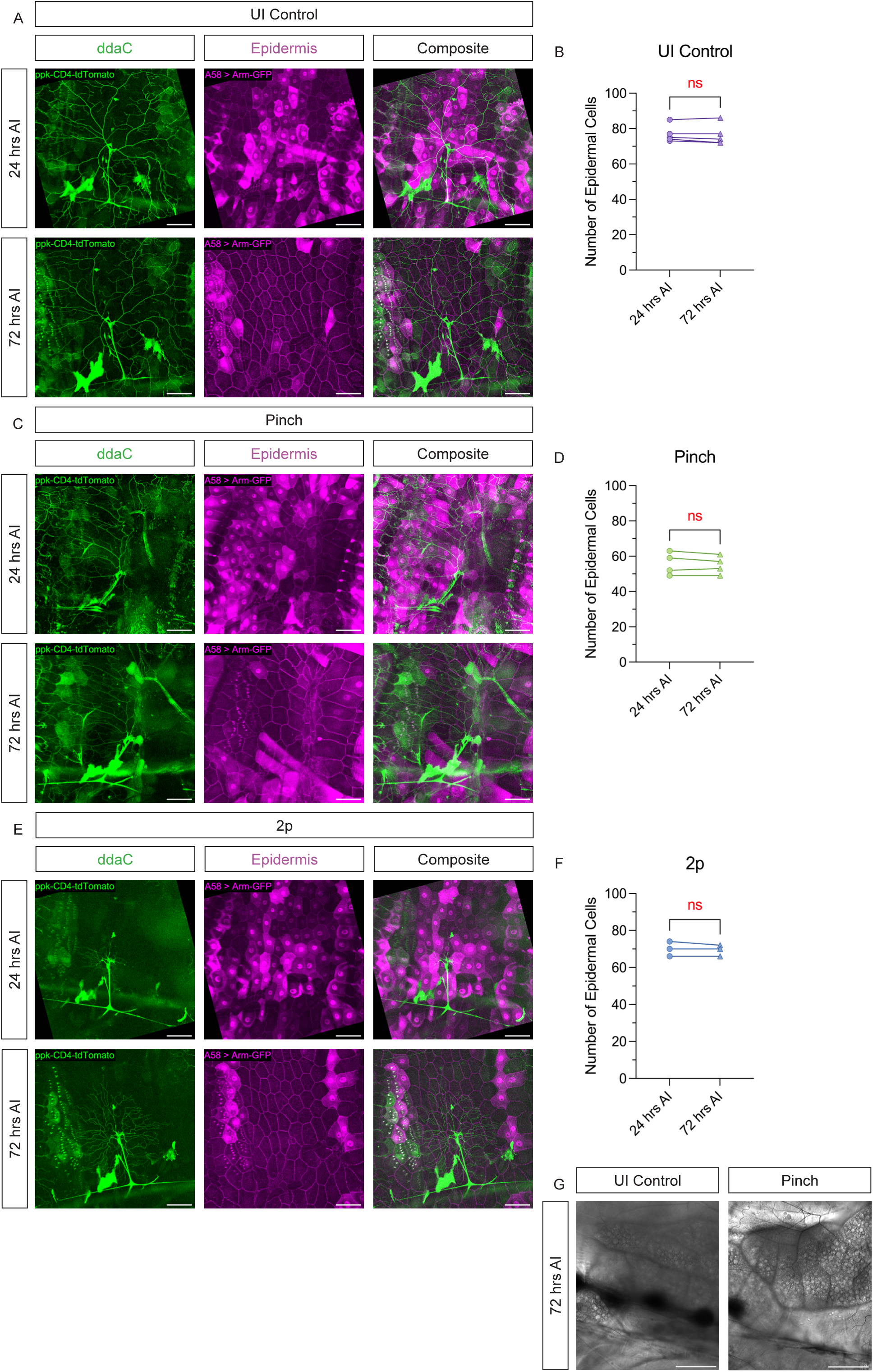
Pinch Injury, but Not 2p-Full Bald Injury, Disrupts Epidermal Cell Morphology. **(A, C, E)** Uninjured (A), pinched (C), or 2p-full bald (E) class IV ddaC neurons with epidermal marker (Gal4^A58^ > UAS-Armadillo-GFP) at 24 and 72 hrs AI. Composite images of pinched or 2p-full bald neurons at 72 hrs AI are also shown in Fig 4E. Scale bar 100 µm. **(B, D, F)** Number of epidermal cells at 24 and 72 hrs AI near uninjured (B), pinched (D), or 2p-full bald (F) neurons. Comparison between injury methods at each time point is also shown in Fig 4F. **(G)** Brightfield images of the cuticle above an uninjured or pinched neuron at 72 hrs AI.

**Figure 4-4:**
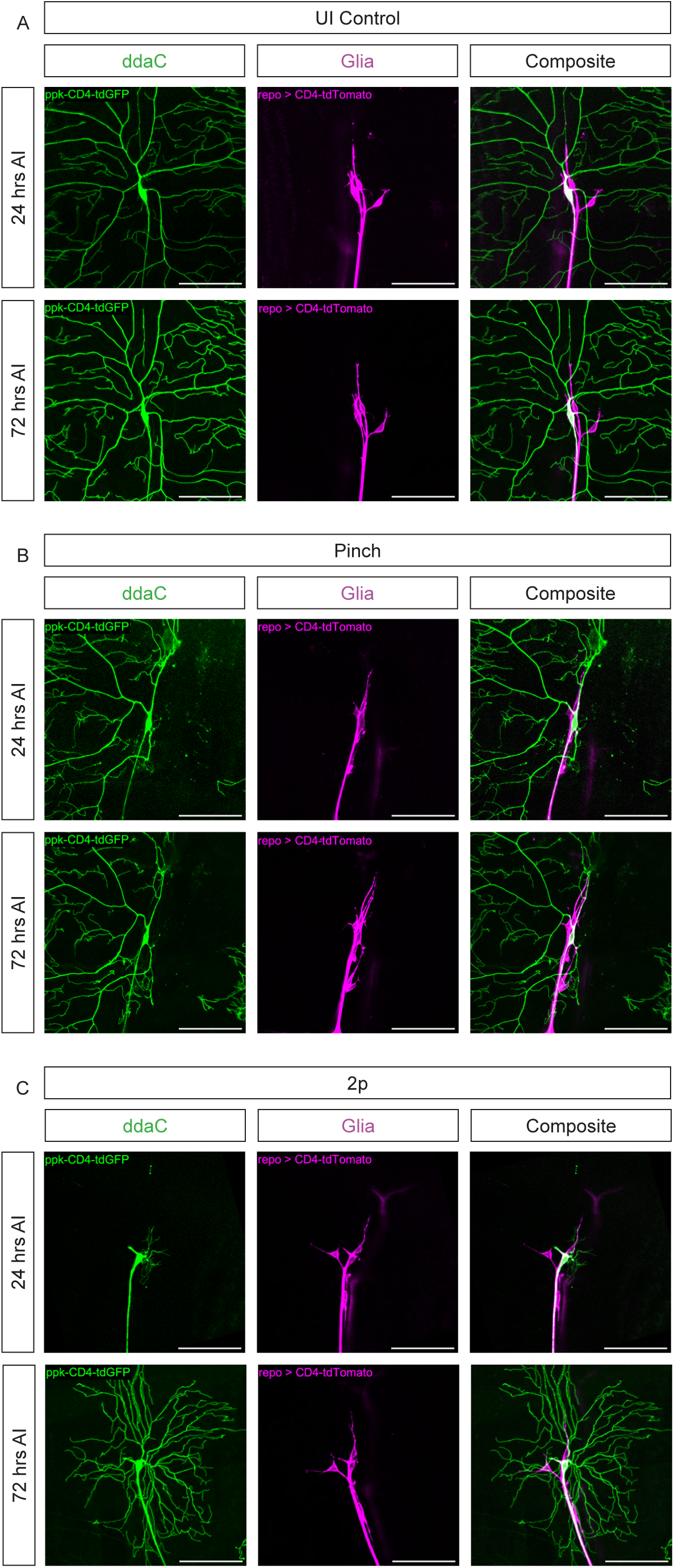
Neither Pinch Nor 2p-Full Bald Injuries Seem to Dramatically Damage Glia. (A-C) Uninjured (A), pinched (B), or 2p-full bald (C) class IV ddaC neurons with glial marker (Gal4^repo^ > CD4-tdTomato) at 24 and 72 hrs AI. Scale bar 100 µm. Composite images of pinched or 2p-full bald neurons at 72 hrs AI are also shown in Fig 4G.

**Figure 4-5:**
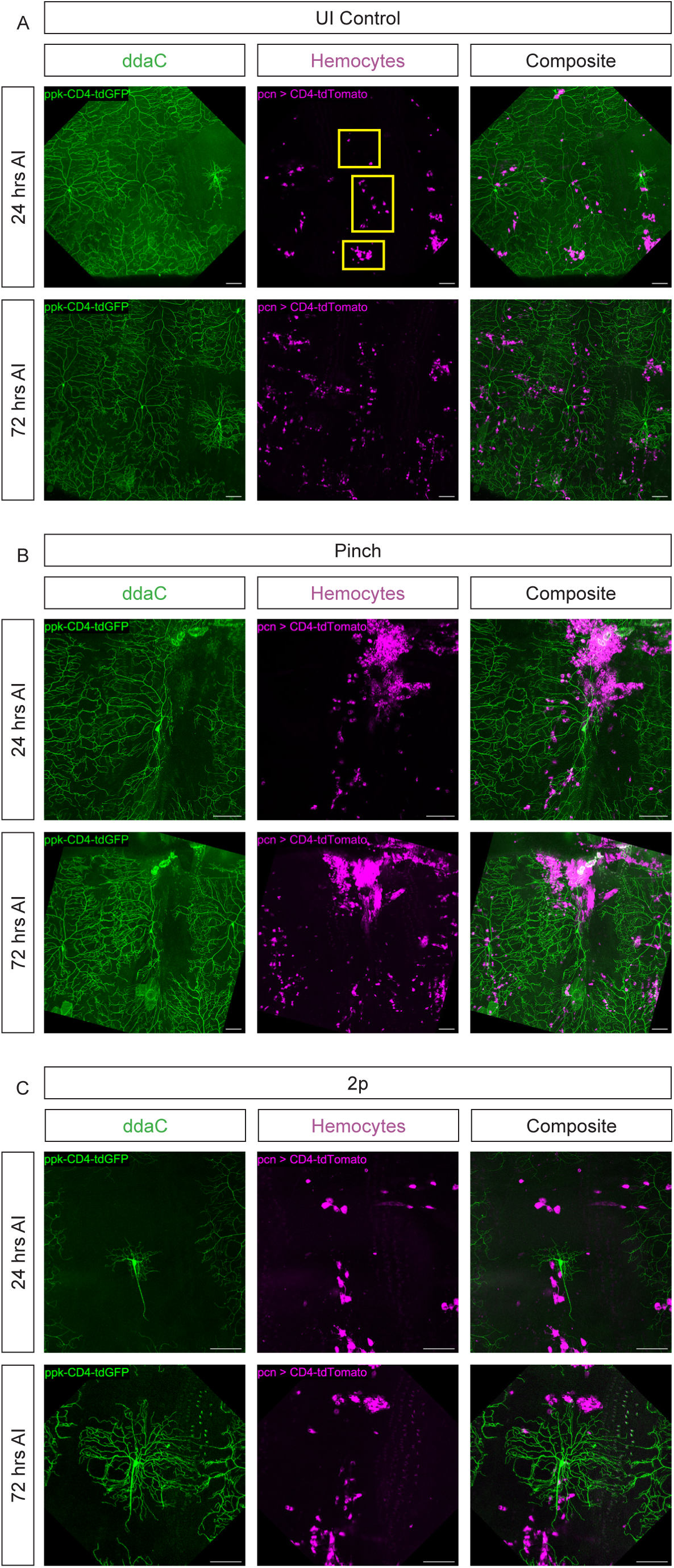
Pinch Injury Recruits Hemocytes. (A-C) Uninjured (A), pinched (B), or 2p-full bald (C) class IV ddaC neurons with hemocyte marker (Gal4^pcn^ > CD4-tdTomato) at 24 and 72 hrs AI. In hemocyte channel at 24 hrs AI, top yellow box identifies dorsal vessel cluster, middle yellow box identifies the dorsal stripe, and the bottom yellow box identifies the lateral patch of hemocytes. Scale bar 100 µm. Composite images of pinched or 2p-full bald neurons at 72 hrs AI are also shown in Fig 4H.

